# Altered co-stimulatory and inhibitory receptors on monocyte subsets in patients with visceral leishmaniasis

**DOI:** 10.1101/2024.03.18.585473

**Authors:** E. Adem, E. Yizengaw, T. Mulaw, E. Nibret, I. Müller, Y. Takele, P. Kropf

**Author notes:** University of Greenwich at Medway, Kent, UK. Department of Comprehensive Cancer Centre, King’s College London, UK. These authors share last authorship (Y. Takele, P. Kropf).

## Abstract

Visceral leishmaniasis (VL) is a neglected tropical disease caused by parasites from the *Leishmania* (*L.*) *donovani* complex. VL is characterised by uncontrolled parasite replication in spleen, liver and bone marrow, and by an impaired immune response and high systemic levels of inflammation. Monocytes have been poorly characterised in VL patients. The aim of this study was to evaluate the expression levels of markers involved in the regulation of T cell responses on different subsets of monocytes from the blood of VL patients and healthy non-endemic controls (HNEC). Monocytes can broadly be divided into three subsets: classical, intermediate and non-classical monocytes. Our results show that the percentages of all three subsets stay similar at the time of VL diagnosis (ToD) and at the end of anti-leishmanial treatment (EoT). We first looked at co-stimulatory receptors: the expression levels of CD40 were significantly increased on classical and intermediate, but not non-classical monocytes, at ToD as compared to EoT and HNEC. CD80 expression levels were also increased on intermediate monocytes at ToD as compared to EoT and HNEC, and on classical monocytes only as compared to HNEC. The levels of CD86 were similar at EoT and ToD and in HNEC on classical and intermediate monocytes, but significantly higher at EoT on non-classical monocytes. We also looked at an inhibitory molecule, PD-L1. Our results show that the expression levels of PD-L1 is significantly higher on all three monocyte subsets at ToD as compared to HNEC, and to EoT on classical and intermediate monocytes.

These results show that monocytes from the blood of VL patients upregulate both co-stimulatory and inhibitory receptors and that their expression levels are restored at EoT.

## INTRODUCTION

Visceral leishmaniasis (VL) is caused by parasites from the *Leishmania donovani* complex, that are transmitted during the blood meal of sand fly vectors. It is endemic in 80 countries, with 12,773 cases reported in 2022 [1]. However, the real number of VL cases are likely to be significantly higher, as the surveillance systems in place are often inadequate and VL occurs in places that are often difficult to access [2, 3]. 90% of VL cases occur in India, Bangladesh, Nepal, Sudan, South Sudan, Ethiopia and Brazil [2].

Whereas the majority of individuals infected with *L. donovani* remain asymptomatic, infection can also develop as a progressive disease. Common symptoms are fever, weight loss, hepatosplenomegaly and pancytopenia [4]. VL patients require hospitalisation and treatment. Existing treatments can cause severe side effects and have long durations. Drug resistance is also a growing problem [5]. One of the primary immunological features of VL patients is their profound immunosuppression: typically, whole blood cells and peripheral blood mononuclear cells (PBMCs) from VL patients display a reduced ability to produce IFNγ and proliferate when exposed to *Leishmania* antigen; and the leishmanin skin test does not induce a delayed-type hypersensitivity reaction. These impaired responses to antigen challenge improve after successful chemotherapy (reviewed in [6–8].

Monocytes in VL patients have been poorly characterised. Monocytes are heterogeneous and can be divided into at least three subsets, based on the expression levels of CD14 and CD16: CD14^high^CD16^low^ (classical), CD14^high^CD16^high^ (intermediate) and CD14^low^CD16^high^ (non-classical) [9, 10]. It has been shown that classical monocytes emerge first in circulation and gradually change their expression levels of CD14 and CD16 into first intermediate monocytes and finally into non-classical monocytes [11]. These three subsets can also each display different effector functions: some of the main functions of classical monocytes are the ability to phagocytose, migrate and produce anti-microbial responses; intermediate monocytes specialise in antigen presentation, regulation of apoptosis and transendothelial migration; and non-classical monocytes into complement and FcR-mediated phagocytosis, transendothelial migration and adhesion (summarised in [12]).

Initial studies showed that monocytes can phagocytose and kill intracellular *L. donovani* by oxidative burst [13–15]. Hoover *et al.* showed that *L. donovani* can replicate within monocytes and that activation of these monocytes with IFNγ results in killing of the parasites [16]; this can be counteracted by IL-4 [17]. Monocytes infected with *L. donovani* did not produce TNFα and IL-1, unless primed with *staphylococcus aureus* or lipopolysaccharide [18]. Monocytes pre-treated with lipophosphoglycan (LPG), a glycolipid present on the surface of parasites, display a lower oxidative burst [19], showing that this protein plays a crucial role in the survival of *Leishmania* parasites in monocytes.

Monocytes from VL patients express lower levels of HLA-DR [20]; as well as CD54 and CD86 and display an impaired oxidative burst [21]. Singh *et al.* showed that monocytes isolated from VL patients exhibit an anti-inflammatory phenotype, characterised by decreased parasite phagocytosis and impaired oxidative burst [22]. We have recently shown that PD-L1 is increased on all three subsets of monocytes at the time of VL diagnosis [23]. In the current study, we extend these results to the analyses of other molecules also involved in the modulation of T cell responses, such as CD40, CD80 and CD86. We assessed their expression levels on the three different subsets of monocytes at the time of VL diagnosis (ToD) and the end of treatment (EoT). These parameters were compared with those of healthy non-endemic controls.

## MATERIALS AND METHODS

### Subjects and sample collection

The study was approved by the Institutional Review Board of the University of Gondar (IRB, reference O/V/P/RCS/05/1572/2017). For this cross-sectional study, 20 male VL patients were recruited at the time of diagnosis (ToD) from the Leishmaniasis Treatment and Research Centre of the Gondar University Hospital; a further 20 males were recruited at the end of successful treatment (EoT), as defined by patients looking improved, afebrile, and having smaller spleen and liver sizes than on admission [24]. 10 non-endemic male controls with no prior history of VL were recruited amongst the staff of the University of Gondar. The median ages of the three cohorts were similar: VL patients at ToD: 25 [20–30]; VL patients at EoT: 23.5 [18.5-26.8] and controls: 24 [20.5-27.3] years old (p=0.7495). Informed written consent was obtained from each patient and control. The exclusion criteria were age (<18 years) and co-infection with HIV. The diagnosis of VL was based on positive serology (rK39) and the presence of amastigotes in spleen or bone marrow aspirates [25]. Patients were treated with a combination of sodium stibogluconate (SSG, 20mg/kg body weight/day) and paromomycin (PM, 15mg/kg body weight/day) injections, given intramuscularly for 17 days.

Ten ml of blood was collected in heparin tubes and was processed immediately after collection: following density gradient centrifugation on Histopaque-1077 (Sigma), the peripheral mononuclear cells (PBMCs) were isolated from the interphase and were used immediately for flowcytometry.

### Flow cytometry

The different subsets of monocytes were defined based on the expression levels of CD14 and CD16 (Figure S1). The following antibodies were used: anti-CD14^APC^ (clone 61D3, eBioscience), anti-CD16^PE^ (clone B73.1, eBioscience) and anti-CD16^FITC^ (clone B73.1, Biolegend). The following antibodies were used to assess the levels of CD40: anti-CD40^PE-Cyanine7^ (clone 5C3, eBioscience); CD80: anti-CD80^FITC^ (clone 2D10.4, eBioscience); CD86: anti-CD86^pe-cyanine7^(clone IT2.2, eBioscience); and PD-L1: anti-PD-L1^PE^ (clone MIH1, eBioscience). The percentages for the isotype controls were <1%. Acquisition was performed using a BD Accuri C6 flow cytometer, at least 5,000 monocytes were acquired, and data were analysed using BD Accuri C6 analysis software.

### Statistical analysis

Data were evaluated for statistical differences using a Kruskal-Wallis test (GraphPad Prism 10) differences were considered statistically significant at *p*<0.05. Unless otherwise specified, results are expressed as median ± interquartile range.

## RESULTS

### Percentages of monocyte subsets

We first determined whether the percentages of the three subsets of monocytes: classical (CD14^high^ CD16^low^), intermediate (CD14^high^ CD16^high^) and non-classical (CD14^low^ CD16^high^) differ between VL patients at the time of diagnosis (ToD) and at the end of treatment (EoT) and as compared to healthy non-endemic controls (HNEC). As expected, the percentages of classical monocytes were higher as compared to the intermediate and the non-classical subsets (Figure 1, Table 1). There were no differences between the percentages of the different subsets at ToD and EoT (Figure 1 and Table 1); as well as in the percentages of classical monocytes at ToD in VL patients and controls, however, there were significantly less classical and more intermediate monocytes in the PBMCs of VL patients at EoT as compared to HNEC (Figure 1, Table 1)).

**Figure 1:**
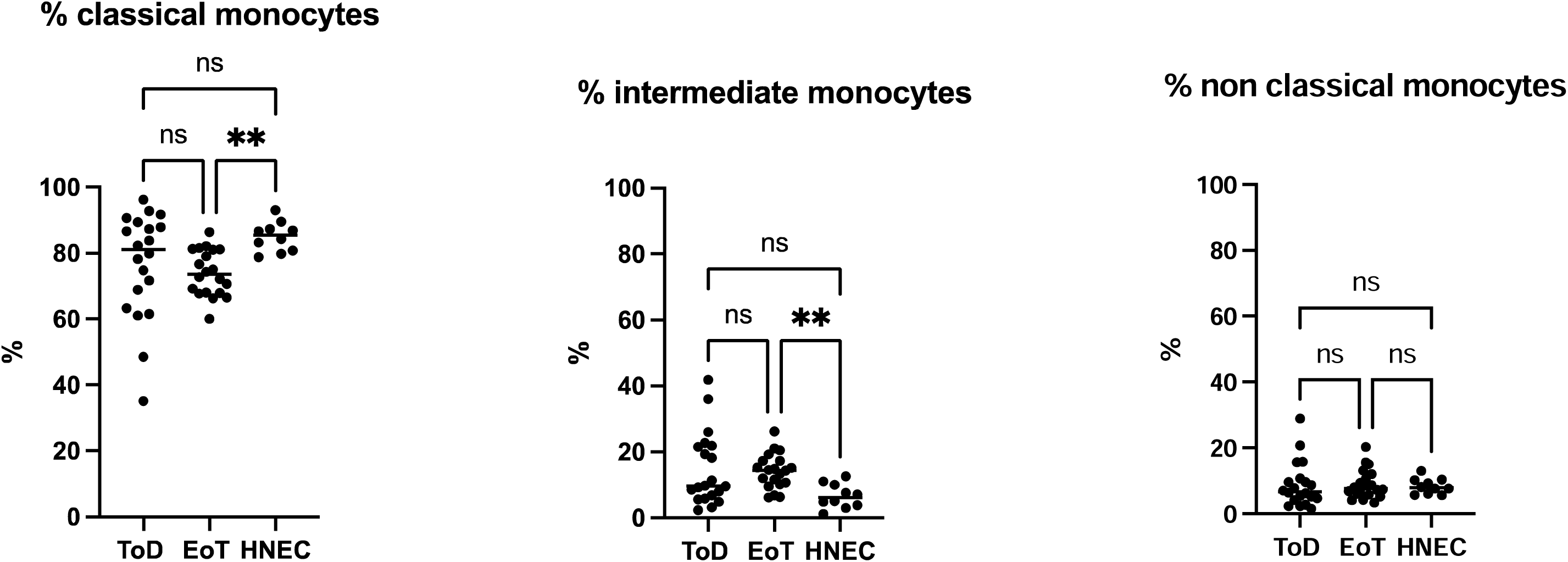
percentages of the different monocyte subsets. PBMCs were purified from VL patients at ToD (n=20) and EoT (n=20) and from HNEC (n=10) as described in Materials and Methods. The percentages of each subset were determined by flow cytometry, as described in Figure S1. Each symbol represents the value for one individual and the straight line represents the median. Statistical differences shown on this figure between VL patients at ToD and EoT and HNEC were determined by Dunn’s multiple comparisons test. ToD=Time of Diagnosis; EoT=End of Treatment; HNEC=healthy non-endemic controls.

**Table 1:**
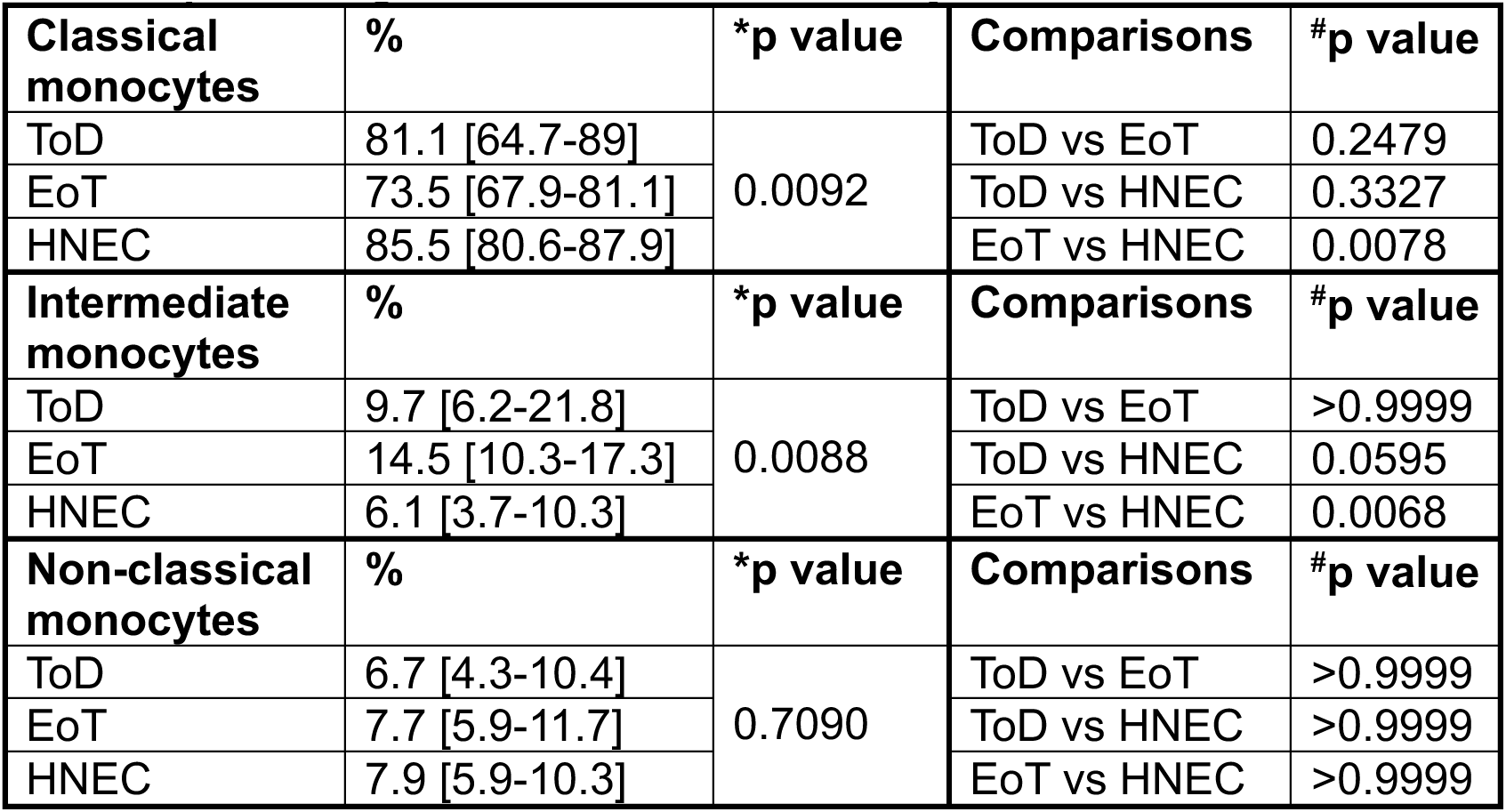
percentages of the different monocyte subsets. PBMCs were purified from VL patients at ToD (n=20) and EoT (n=20) and from HNEC (n=10) and the percentages of each subset were determined by flow cytometry, as described in Figure S1. Results are presented as median with interquartile range. Statistical differences were determined by Kruskal-Wallis test (*) and Dunn’s multiple comparisons test (^#^). ToD=Time of Diagnosis; EoT=End of Treatment; HNEC=healthy non-endemic controls.

| **Classical monocytes** | **%** | **\*p value** | **Comparisons** | **#p value** |
| --- | --- | --- | --- | --- |
| ToD | 81.1 [64.7-89] | 0.0092 | ToD vs EoT | 0.2479 |
| EoT | 73.5 [67.9-81.1] |  | ToD vs HNEC | 0.3327 |
| HNEC | 85.5 [80.6-87.9] |  | EoT vs HNEC | 0.0078 |
| **Intermediate monocytes** | **%** | **\*p value** | **Comparisons** | **#p value** |
| ToD | 9.7 [6.2-21.8] | 0.0088 | ToD vs EoT | >0.9999 |
| EoT | 14.5 [10.3-17.3] |  | ToD vs HNEC | 0.0595 |
| HNEC | 6.1 [3.7-10.3] |  | EoT vs HNEC | 0.0068 |
| **Non-classical monocytes** | **%** | **\*p value** | **Comparisons** | **#p value** |
| ToD | 6.7 [4.3-10.4] | 0.7090 | ToD vs EoT | >0.9999 |
| EoT | 7.7 [5.9-11.7] |  | ToD vs HNEC | >0.9999 |
| HNEC | 7.9 [5.9-10.3] |  | EoT vs HNEC | >0.9999 |

### Activation status of the different monocyte subsets

We first measured the expression levels of CD14 and CD16. Results presented in Table 2 show no significant differences between the different subsets in VL patients at ToD and EoT and in HNEC.

**Table 2:** MFI of CD14 and CD16 on the different monocyte subsets. PBMCs were purified from VL patients at ToD (n=20) and EoT (n=20) and from HNEC (n=10) and the expression levels (MFI = Median Fluorescence Intensity) of CD14 and CD16 were measured on the different monocyte subsets by flow cytometry. Results are presented as median with interquartile range. Statistical differences were determined by Kruskal-Wallis test (*). ToD=Time of Diagnosis; EoT=End of Treatment; HNEC=healthy non-endemic controls.

| **Classical monocytes** | **MFI CD14 (x10<sup>3</sup>)** | **\*p value** | **MFI CD16 (x10<sup>3</sup>)** | **\*p value** |
| --- | --- | --- | --- | --- |
| ToD | 15.2 [10.6-15.2] | 0.0823 | 0.9 [0.8-1.3] | 0.1180 |
| EoT | 17.1 [13.1-20.8] |  | 0.9 [0.7-1.3] |  |
| HNEC | 21.3 [16-23.1] |  | 0.6 [0.51-1.3] |  |
| **Intermediate monocytes** | **MFI CD14 (x10<sup>3</sup>)** | **\*p value** | **MFI CD16 (x10<sup>3</sup>)** | **\*p value** |
| ToD | 14 [10.9-2.2] | 0.4088 | 10.2 [8.5-13.1] | 0.5551 |
| EoT | 16.2 [11.8-19.1] |  | 10 [7-11.9] |  |
| HNEC | 16.7 [13.3-23.5] |  | 8 [6.1-28.2] |  |
| **Non-classical monocytes** | **MFI CD14 (x10<sup>3</sup>)** | **\*p value** | **MFI CD16 (x10<sup>3</sup>)** | **\*p value** |
| ToD | 1.3 [0.4-2.9] | 0.3888 | 16.5 [13.5-34.6] | 0.3818 |
| EoT | 1.9 [1.6-2.5] |  | 19.8 [15.5-36.9] |  |
| HNEC | 2.1 [1.3-3.2] |  | 27.2 [20.7-42.6] |  |

Next, we measured the expression levels of the co-stimulatory molecules CD40, CD80 and CD86 and the immune regulatory molecule PD-L1 on the different monocyte subsets.

Results presented in Figure 2 and Table 3 show that CD40 MFI was significantly higher at ToD as compared to EoT and as compared to HNEC on the classical and intermediate monocyte subsets isolated from VL patients.

**Figure 2:**
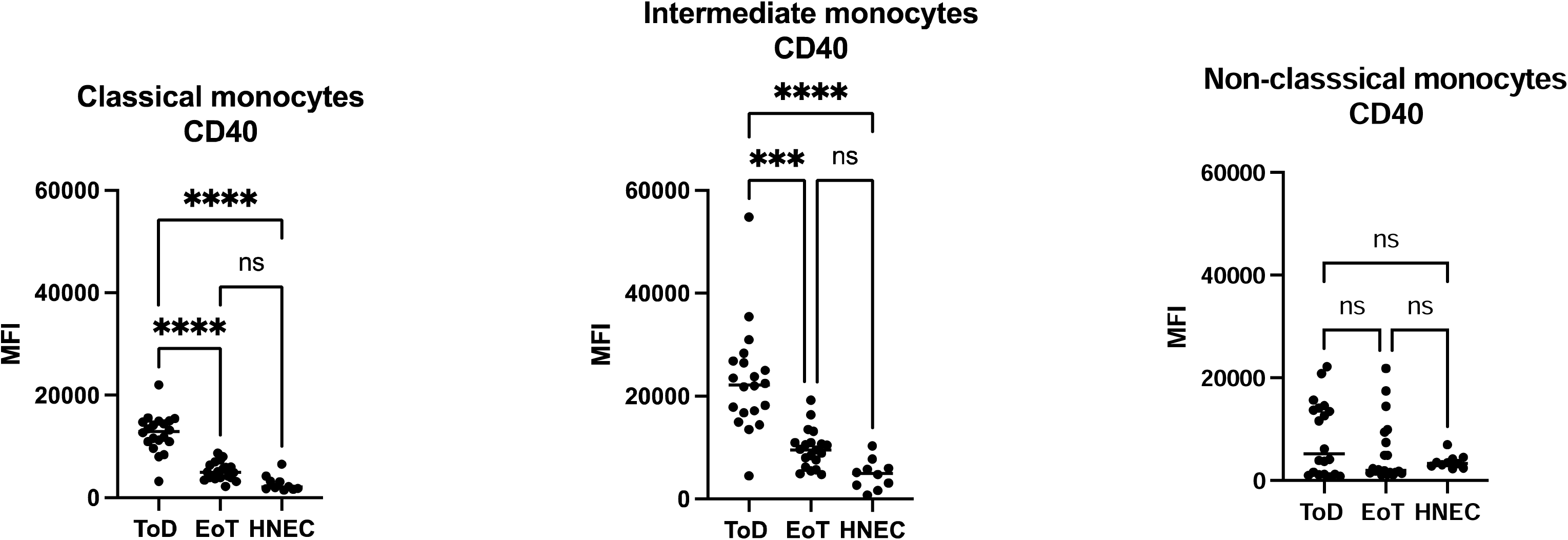
Expression levels of CD40 on the different monocyte subsets. PBMCs were purified from VL patients at ToD (n=20) and EoT (n=20) and from HNEC (n=10) as described in Materials and Methods. The expression levels (MFI = Median Fluorescence Intensity) of CD40 were measured on the different monocyte subsets by flow cytometry as described in Materials and Methods. Each symbol represents the value for one individual and the straight line represents the median. Statistical differences shown on this figure between VL patients at ToD and EoT and HNEC were determined by Dunn’s multiple comparisons test. ToD=Time of Diagnosis; EoT=End of Treatment; HNEC=healthy non-endemic controls.

**Table 3:** CD40 MFI on the different monocyte subsets. PBMCs were purified from VL patients at ToD (n=20) and EoT (n=20) and from HNEC (n=10) and the expression levels (MFI = Median Fluorescence Intensity) of CD40 were measured on the different monocyte subsets by flow cytometry. Results are presented as median with interquartile range. Statistical differences were determined by Kruskal-Wallis test (*) and Dunn’s multiple comparisons test (^#^). ToD=Time of Diagnosis; EoT=End of Treatment; HNEC=healthy non-endemic controls.

| **Classical monocytes** | **CD40 MFI (x10<sup>3</sup>)** | **\*p value** | **Comparisons CD40 MFI** | **#p value** |
| --- | --- | --- | --- | --- |
| ToD | 13 [11-14.9] | <0.0001 | ToD vs EoT | <0.0001 |
| EoT | 5 [4-6.3] |  | ToD vs HNEC | <0.0001 |
| HNEC | 2.1 [1.7-3.5] |  | EoT vs HNEC | 0.1236 |
| **Intermediate monocytes** | **%** | **\*p value** | **Comparisons CD40 MFI** | **#p value** |
| ToD | 22.2 [16.9-26.7] | <0.0001 | ToD vs EoT | 0.0003 |
| EoT | 9.6 [6.6-11] |  | ToD vs HNEC | <0.0001 |
| HNEC | 5 [2.5-6.4] |  | EoT vs HNEC | 0.1063 |
| **Non-classical monocytes** | **%** | **\*p value** | **Comparisons CD40 MFI** | **#p value** |
| ToD | 5.2 [1.2-14] | 0.6399 | ToD vs EoT | >0.9999 |
| EoT | 2.0 [1.5-8.9] |  | ToD vs HNEC | >0.9999 |
| HNEC | 3.4 [2.7-4.3] |  | EoT vs HNEC | >0.9999 |

There were no significant differences for these 2 subsets between VL patients at EoT and HNEC; or between VL patients at ToD and EoT and HNEC on non-classical monocytes (Figure 2 and Table 3). Of note, the intermediate monocytes from VL patients expressed the highest levels of CD40 as compared to the classical and the non-classical monocytes at both time points. The MFI of CD40 was similar on all three subsets of monocytes from HNEC (Table S1).

CD80 MFI was significantly higher on classical and intermediate monocytes from VL patients at ToD as compared to HNEC and between ToD and EoT on intermediate monocytes. There were no significant differences between VL patients at EoT and HNEC; or between VL patients at ToD and EoT and HNEC on non-classical monocytes (Figure 3 and Table 4). The classical monocytes from VL patients at EoT and ToD and from the HNEC expressed the lowest levels of CD80 (Table S2).

**Figure 3:**
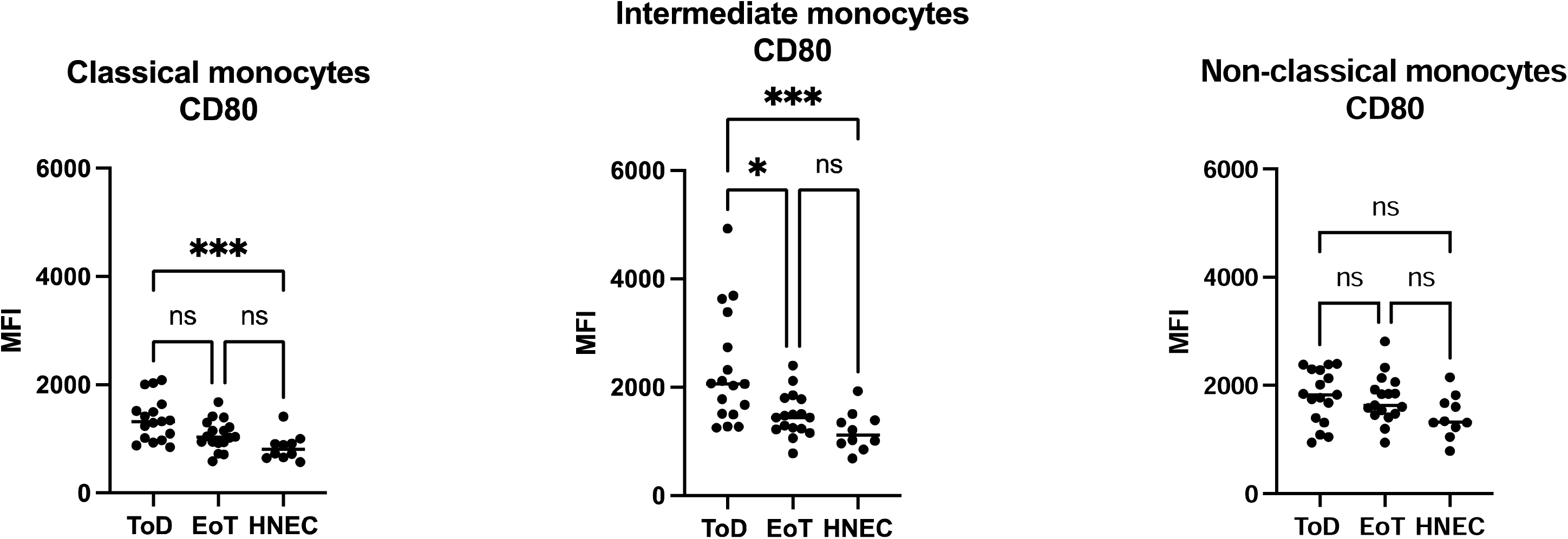
Expression levels of CD80 on the different monocyte subsets. PBMCs were purified from VL patients at ToD (n=17) and EoT (n=17) and from HNEC (n=10) as described in Materials and Methods. The expression levels (MFI = Median Fluorescence Intensity) of CD80 were measured on the different monocyte subsets by flow cytometry as described in Materials and Methods. Each symbol represents the value for one individual and the straight line represents the median. Statistical differences shown on this figure between VL patients at ToD and EoT and HNEC were determined by Dunn’s multiple comparisons test. ToD=Time of Diagnosis; EoT=End of Treatment; HNEC=healthy non-endemic controls.

**Table 4:** CD80 MFI on the different monocyte subsets. PBMCs were purified from VL patients at ToD (n=17) and EoT (n=17) and from HNEC (n=10) and the expression levels (MFI = Median Fluorescence Intensity) of CD80 were measured on the different monocyte subsets by flow cytometry. Results are presented as median with interquartile range. Statistical differences were determined by Kruskal-Wallis test (*) and Dunn’s multiple comparisons test (^#^). ToD=Time of Diagnosis; EoT=End of Treatment; HNEC=healthy non-endemic controls.

| **Classical monocytes** | **CD80 MFI** | **\*p value** | **Comparisons CD80 MFI** | **#p value** |
| --- | --- | --- | --- | --- |
| ToD | 1321 [998-1579] | 0.0014 | ToD vs EoT | 0.2240 |
| EoT | 1039 [930-1260] |  | ToD vs HNEC | 0.0009 |
| HNEC | 807 [657-934] |  | EoT vs HNEC | 0.1134 |
| **Intermediate monocytes** | **CD80 MFI** | **\*p value** | **Comparisons CD80 MFI** | **#p value** |
| ToD | 2060 [1506-3062] | 0.0005 | ToD vs EoT | 0.0311 |
| EoT | 1445 [1231-1791] |  | ToD vs HNEC | 0.0005 |
| HNEC | 1120 [937-1420] |  | EoT vs HNEC | 0.3447 |
| **Non-classical monocytes** | **CD80 MFI** | **\*p value** | **Comparisons CD80 MFI** | **#p value** |
| ToD | 1826 [1356-2289] | 0.0947 | ToD vs EoT | >0.9999 |
| EoT | 1630 [1464-1993] |  | ToD vs HNEC | 0.1058 |
| HNEC | 1325 [1183-1706] |  | EoT vs HNEC | 0.2469 |

CD86 was significantly lower on non-classical monocytes at ToD as compared to EoT. All the other comparisons were not significant (Figure 4 and Table 5). Whereas intermediate and non-classical monocytes from HNEC and VL patients at EoT expressed higher levels of CD86 as compared to the classical monocytes (Table S3), at ToD, the intermediate subsets expressed the highest levels of CD86 (Table S3).

**Figure 4:**
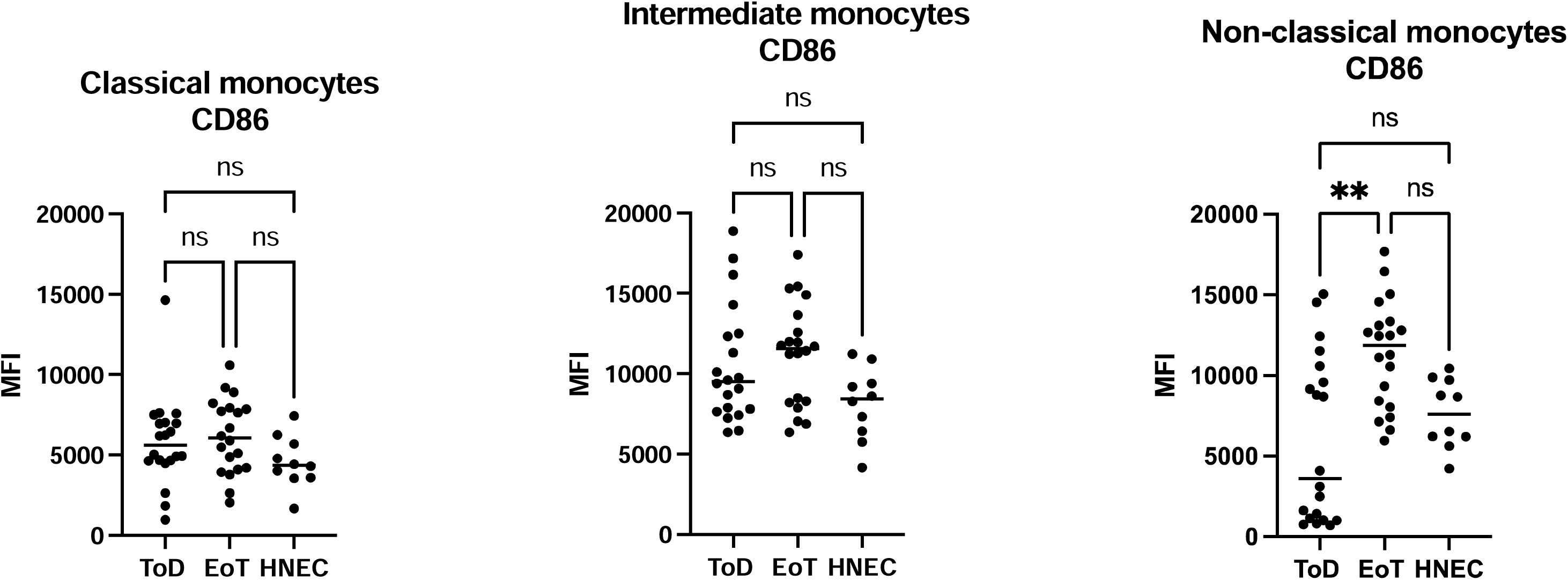
Expression levels of CD86 on the different monocyte subsets. PBMCs were purified from VL patients at ToD (n=20) and EoT (n=20) and from HNEC (n=10) as described in Materials and Methods. The expression levels (MFI = Median Fluorescence Intensity) of CD86 were measured on the different monocyte subsets by flow cytometry as described in Materials and Methods. Each symbol represents the value for one individual and the straight line represents the median. Statistical differences shown on this figure between VL patients at ToD and EoT and HNEC were determined by Dunn’s multiple comparisons test. ToD=Time of Diagnosis; EoT=End of Treatment; HNEC=healthy non-endemic controls.

**Table 5:** CD86 MFI on the different monocyte subsets. PBMCs were purified from VL patients at ToD (n=20) and EoT (n=20) and from HNEC (n=10) and the expression levels (MFI = Median Fluorescence Intensity) of CD86 were measured on the different monocyte subsets by flow cytometry. Results are presented as median with interquartile range. Statistical differences were determined by Kruskal-Wallis test (*) and Dunn’s multiple comparisons test (^#^). ToD=Time of Diagnosis; EoT=End of Treatment; HNEC=healthy non-endemic controls.

| **Classical monocytes** | **CD86 MFI** | **\*p value** | **Comparisons** | **#p value** |
| --- | --- | --- | --- | --- |
|  |  |  | **CD86 MFI** |  |
| ToD | 5606 [4646-7007] | 0.1158 | ToD vs EoT | >0.9999 |
| EoT | 6051 [4133-7919] |  | ToD vs HNEC | 0.3864 |
| HNEC | 4374 [3587-5841] |  | EoT vs HNEC | 0.1159 |
| **Intermediate monocytes** | **CD86 MFI** | **\*p value** | **Comparisons** | **#p value** |
|  |  |  | **CD86 MFI** |  |
| ToD | 9500 [7688-12461] | 0.0585 | ToD vs EoT | >0.9999 |
| EoT | 11569 [8234-13889] |  | ToD vs HNEC | 0.2697 |
| HNEC | 8436 [6258-9775] |  | EoT vs HNEC | 0.0522 |
| **Non-classical monocytes** | **CD86 MFI** | **\*p value** | **Comparisons** | **#p value** |
|  |  |  | **CD86 MFI** |  |
| ToD | 3604 [1054-10332] | 0.0018 | ToD vs EoT | 0.0020 |
| EoT | 11877 [8136-13303] |  | ToD vs HNEC | >0.9999 |
| HNEC | 7600 [6065-9769] |  | EoT vs HNEC | 0.0646 |

**Table 6:** PDL-1 MFI on the different monocyte subsets. PBMCs were purified from VL patients at ToD (n=20) and EoT (n=20) and from HNEC (n=10) and the expression levels (MFI = Median Fluorescence Intensity) of PD-L1 were measured on the different monocyte subsets by flow cytometry. Results are presented as median with interquartile range. Statistical differences were determined by Kruskal-Wallis test (*) and Dunn’s multiple comparisons test (^#^). ToD=Time of Diagnosis; EoT=End of Treatment; HNEC=healthy non-endemic controls.

| **Classical monocytes** | **PDL-1 MFI** | **\*p value** | **Comparisons** | **#p value** |
| --- | --- | --- | --- | --- |
|  |  |  | **PDL-1 MFI** |  |
| ToD | 2951 [2132-4051] | <0.0001 | ToD vs EoT | 0.0052 |
| EoT | 1233 [1009-2076] |  | ToD vs HNEC | <0.0001 |
| HNEC | 893 [687-1205] |  | EoT vs HNEC | 0.1525 |
| **Intermediate monocytes** | **PDL-1 MFI** | **\*p value** | **Comparisons** | **#p value** |
|  |  |  | **PDL-1 MFI** |  |
| ToD | 5779 [4578-8411] | <0.0001 | ToD vs EoT | 0.0005 |
| EoT | 2058 [1656-3169] |  | ToD vs HNEC | <0.0001 |
| HNEC | 1269 [996-2233] |  | EoT vs HNEC | 0.2273 |
| **Non-classical monocytes** | **PDL-1 MFI** | **\*p value** | **Comparisons** | **#p value** |
|  |  |  | **PDL-1 MFI** |  |
| ToD | 3451 [1183-4191] | 0.0096 | ToD vs EoT | 0.4566 |
| EoT | 1923 [1365-2506] |  | ToD vs HNEC | 0.0070 |
| HNEC | 1212 [887-1604] |  | EoT vs HNEC | 0.1832 |

Results presented in Figure 5 show that at ToD, PD-L1 expression levels on all three different subsets were significantly higher as compared to those from HNEC. PDL-1 MFI decreased significantly at EoT on classical and intermediate, but not on non-classical monocytes. There were no significant differences in PD-L1 MFI between EoT and HNEC in all three monocyte subsets. Whereas intermediate and non-classical monocytes from HNEC and VL patients at EoT expressed higher levels of PD-L1 as compared to the classical monocytes (Table S4), at ToD, the intermediate expressed the highest levels of PD-L1 (Table S4).

**Figure 5:**
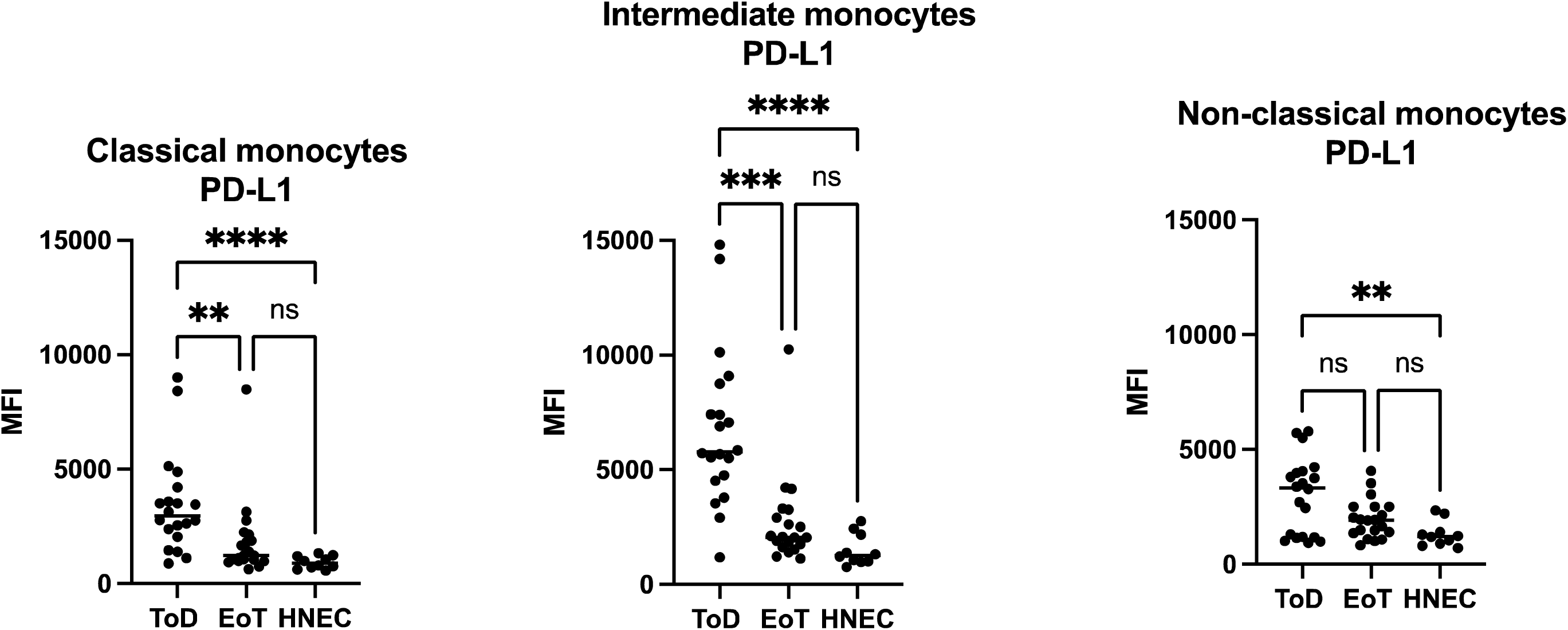
Expression levels of PD-L1 on the different monocyte subsets. PBMCs were purified from VL patients at ToD (n=20) and EoT (n=20) and from HNEC (n=10) as described in Materials and Methods. The expression levels (MFI = Median Fluorescence Intensity) of PD-L1 were measured on the different monocyte subsets by flow cytometry as described in Materials and Methods. Each symbol represents the value for one individual and the straight line represents the median. Statistical differences shown on this figure between VL patients at ToD and EoT and HNEC were determined by Dunn’s multiple comparisons test. ToD=Time of Diagnosis; EoT=End of Treatment; HNEC=healthy non-endemic controls.

## DISCUSSION

Our results show that the percentages of classical, intermediate and non-classical monocytes in the blood of VL patients are similar at ToD and EoT. However, the expression levels of CD40, CD80, CD86 and PD-L1 vary on the different monocyte subsets and some of these markers change their expression levels at the end of anti-leishmanial treatment. Our results also show that at EoT, all of these markers are back to similar levels as those from HNEC.

CD40 is a member of the tumor necrosis factor (TNF) receptor family and is expressed on B cells, monocytes, dendritic cells, macrophages as well as on non-hematopoietic cells such as fibroblast (summarised in [26]). Its ligand, CD40L, is predominantly expressed on activated T and B cells and on platelets and can be shed (soluble CD40L (sCD40L)). CD40/CD40L interaction stimulates cells to produce chemokines and cytokines, and upregulate co-stimulatory and adhesion molecules, as well as enzymes, such as matrix metalloproteinases [27, 28].

Little is known about CD40 expression on monocytes in VL patients. The levels of the soluble form of its ligand, sCD40L, have been shown to be lower in plasma of HIV/VL patients as compared to patients with HIV asymptomatically co-infected with *Leishmania* parasites [29]. De Oliveira *et al*. showed that *Leishmania infantum*-infected macrophages exposed to sCD40L harbour less parasites [30]. High levels of sCD40L in serum are associated with successful anti-leishmanial treatment [31]. Here we show that CD40 MFI is increased on both classical and intermediate monocytes at ToD and decreased at EoT. CD40 is expressed at low levels on monocytes and its upregulation is associated to inflammatory conditions such as atherosclerosis [32], multiple sclerosis [33] and tuberculosis [34]. It is therefore possible that the increased CD40 MFI measured on monocytes isolated from VL patients at ToD is a result of the high level of inflammatory cytokines we measured in the plasma of these patients [35]. The levels of CD40 MFI could also be increased as a positive feedback loop to boost T cell responses, as these are inefficient during the active phase of VL [35]. At EoT, CD40 expression levels decreased, concomitantly with a partial restoration of T cell response and lower levels of inflammatory cytokines [35]. Since CD40 was upregulated on classical and intermediate, but not on non-classical monocytes, these results suggest that the latter subset might not be as important in T cell activation via CD40.

CD80 and CD86 bind to two receptors on the surface of T cells: CD28 and cytotoxic T-lymphocyte associated protein 4 (CTLA-4). The interaction with CD28 results in T cell activation and that with CTLA-4 in T cell suppression [36]. CD80 binds more efficiently to CTLA-4 as compared to CD28, and CD86 binds to both receptors less efficiently than CD80 [37]. It has also been shown that CD80 can interact with PD-L1 and thereby prevent the binding of PD-1 with PD-L1 [38]. Here we show that the expression levels of CD80 were higher at ToD on classical and intermediate monocytes. Since CD80 has a higher affinity to CTLA-4, this might contribute to the impaired antigen-specific responses observed at ToD [35]. CD86 was significantly higher on non-classical monocytes at EoT as compared to ToD. This suggests that non-classical monocyte might play a role in T cell activation via CD86 at ToD.

We have recently shown that PD-L1 was increased on all three subsets of monocytes at ToD [23]. Here we extended these results to EoT and show that PD-L1 was significantly decreased on classical and intermediate monocytes. This concurs with the more efficient immune response observed at EoT in VL patients, and indeed, our previous results show that PD1 was also lower on CD4+ T cells at EoT [35].

In summary our results show that monocytes upregulate both co-stimulatory and inhibitory molecules during VL and that at EoT, these levels are similar to those of HNEC. We haven’t measured the levels of CD28, CD40L and CTLA-4 on T cells at ToD and EoT. All these molecules exist as a soluble form and could interact with their receptors on monocytes: for example, sCTLA-4 could interact with CD80/86 and thereby prevent the binding with CD28 and T cell activation, as shown in autoimmune diseases [39]. The interaction of the inhibitory molecules PD-1 and PD-L1 is better characterised in VL: we have recently shown that IFNy production is improved by inhibition of PD-1/PD-L1 ligation [23]. Therefore, the high expression of both molecules at ToD and their lower expression at EoT is in agreement with the efficiency of the immune response at these two time points [35].

Even though monocytes can be broadly divided into classical, intermediate and non-classical subsets based on the expression levels of CD14 and CD16, discrepancies, as well as redundancies in effector functions have been shown between the different subsets (summarised in [10]). Recently, more subsets have been identified with techniques including mass cytometry [40] and by using other markers such as CD62L, CD49d and CD43 and size [41]. Due to the heterogeneity of monocytes, conducting more detailed functional and phenotypic analyses of the different subsets will contribute to a better understanding of the role of these cells in the immunopathology of visceral leishmaniasis.

## ACKNOWLEDGMENTS

We thank Arega Yeshanew, Roma Melkamu, Saba Atnafu, Aschalew Tamiru, Zemenay Mulugeta, Tigist Mekonen, Yohannes Sinku and Eden G/Hiwot from the University of Gondar for their invaluable help and enthusiastic cooperation in this study.

