## Supplementary tables for "Altered co-stimulatory and inhibitory receptors on monocyte subsets in patients with visceral leishmaniasis"

**Table S1: CD40 MFI on monocyte subsets from VL patients at ToD and EoT and on monocytes from HNEC**

| **ToD** | **CD40 MFI ($x10^3$)** | ***p value** | **Comparisons CD40 MFI** | **#p value** |
|---|---|---|---|---|
| Classical | 13 [11-14.9] | <0.0001 | C vs I | 0.0007 |
| Intermediate | 22.2 [16.9-26.7] | | C vs NC | 0.4656 |
| Non-classical | 5.2 [1.2-14] | | I vs NC | <0.0001 |
| **EoT** | **CD40 MFI ($x10^3$)** | ***p value** | **Comparisons CD40 MFI** | **#p value** |
| Classical | 5 [4-6.3] | 0.0001 | C vs I | 0.0034 |
| Intermediate | 9.6 [6.6-11] | | C vs NC | >0.9999 |
| Non-classical | 2.0 [1.5-8 | | I vs NC | 0.0002 |
| **HNEC** | **CD40 MFI ($x10^3$)** | ***p value** | **Comparisons CD40 MFI** | **#p value** |
| Classical | 2.1 [1.7-3.5] | 0.1050 | C vs I | 0.1265 |
| Intermediate | 5 [2.5-6.4] | | C vs NC | 0.3639 |
| Non-classical | 3.4 [2.7-4.3] | | I vs NC | >0.9999 |

PBMCs were purified from VL patients at ToD (n=20) and EoT (n=20) and from HNEC (n=10) and the expression levels (MFI = Median Fluorescence Intensity) of CD40 were measured on the different monocyte subsets by flow cytometry. Results are presented as median with interquartile range. Statistical differences were determined by Kruskall-Wallis test (*) and Dunn's multiple comparisons test (#). ToD=Time of Diagnosis; EoT=End of Treatment; HNEC=healthy non-endemic controls. C=classical monocytes. I=intermediate monocytes. NC=non-classical monocytes.

**Table S2: CD80 MFI on monocyte subsets from VL patients at ToD and EoT and on monocytes from HNEC**

| ToD | CD80 MFI | *p value | Comparisons CD80 MFI | #p value |
|---|---|---|---|---|
| Classical | 1321 [998-1579] | 0.0021 | C vs I | 0.0020 |
| Intermediate | 2060 [1506-3062] | | C vs NC | 0.0434 |
| Non-classical | 1826 [1356-2289] | | I vs NC | >0.9999 |
| **EoT** | **CD80 MFI** | ***p value** | **Comparisons CD80 MFI** | **#p value** |
| Classical | 1039 [930-1260] | <0.0001 | C vs I | 0.0081 |
| Intermediate | 1445 [1231-1791] | | C vs NC | <0.0001 |
| Non-classical | 1630 [1464-1993] | | I vs NC | 0.3922 |
| **HNEC** | **CD80 MFI** | ***p value** | **Comparisons CD80 MFI** | **#p value** |
| Classical | 807 [657-934] | 0.0042 | C vs I | 0.0925 |
| Intermediate | 1120 [937-1420] | | C vs NC | 0.0034 |
| Non-classical | 1325 [1183-1706] | | I vs NC | 0.8242 |

PBMCs were purified from VL patients at ToD (n=17) and EoT (n=17) and from HNEC (n=10) and the expression levels (MFI = Median Fluorescence Intensity) of CD80 were measured on the different monocyte subsets by flow cytometry. Results are presented as median with interquartile range. Statistical differences were determined by Kruskall-Wallis test (*) and Dunn's multiple comparisons test (#). ToD=Time of Diagnosis; EoT=End of Treatment; HNEC=healthy non-endemic controls. C=classical monocytes. I=intermediate monocytes. NC=non-classical monocytes.

**Table S3: CD86MFI on monocyte subsets from VL patients at ToD and EoT and on monocytes from HNEC**

| **ToD** | **CD86 MFI** | ***p value** | **Comparisons CD86 MFI** | **#p value** |
|---|---|---|---|---|
| Classical | 5606 [4646-7007] | | C vs I | 0.0008 |
| Intermediate | 9500 [7688-12461] | 0.0003 | C vs NC | >0.9999 |
| Non-classical | 3604 [1054-10332] | | I vs NC | 0.0036 |
| **EoT** | **CD86 MFI** | ***p value** | **Comparisons CD86 MFI** | **#p value** |
| Classical | 6051 [4133-7919] | | C vs I | <0.0001 |
| Intermediate | 11569 [8234-13889] | <0.0001 | C vs NC | <0.0001 |
| Non-classical | 11877 [8136-13303] | | I vs NC | >0.9999 |
| **HNEC** | **CD86 MFI** | ***p value** | **Comparisons CD86 MFI** | **#p value** |
| Classical | 4374 [3587-5841] | | C vs I | 0.0063 |
| Intermediate | 8436 [6258-9775] | 0.0035 | C vs NC | 0.0197 |
| Non-classical | 7600 [6065-9769] | | I vs NC | >0.9999 |

PBMCs were purified from VL patients at ToD (n=20) and EoT (n=20) and from HNEC (n=10) and the expression levels (MFI = Median Fluorescence Intensity) of CD86 were measured on the different monocyte subsets by flow cytometry. Results are presented as median with interquartile range. Statistical differences were determined by Kruskall-Wallis test (*) and Dunn's multiple comparisons test (#). ToD=Time of Diagnosis; EoT=End of Treatment; HNEC=healthy non-endemic controls. C=classical monocytes. I=intermediate monocytes. NC=non-classical monocytes.

**Table S4: PD-L1 MFI on monocyte subsets from VL patients at ToD and EoT and on monocytes from HNEC**

| ToD | PD-L1 MFI | *p value | Comparisons PD-L1 MFI | #p value |
|---|---|---|---|---|
| Classical | 2951 [2132-4051] | 0.0001 | C vs I | 0.0008 |
| Intermediate | 5779 [4578-8411] | | C vs NC | >0.9999 |
| Non-classical | 3451 [1183-4191] | | I vs NC | 0.0007 |
| **EoT** | **PD-L1 MFI** | ***p value** | **Comparisons PD-L1 MFI** | **#p value** |
| Classical | 1233 [1009-2076] | 0.0131 | C vs I | 0.0099 |
| Intermediate | 2058 [1656-3169] | | C vs NC | 0.3038 |
| Non-classical | 1923 [1365-2506] | | I vs NC | 0.5816 |
| **HNEC** | **PD-L1 MFI** | ***p value** | **Comparisons PD-L1 MFI** | **#p value** |
| Classical | 893 [687-1205] | 0.0572 | C vs I | 0.0668 |
| Intermediate | 1269 [996-2233] | | C vs NC | 0.2390 |
| Non-classical | 1212 [887-1604] | | I vs NC | >0.9999 |

PBMCs were purified from VL patients at ToD (n=20) and EoT (n=20) and from HNEC (n=10) and the expression levels (MFI = Median Fluorescence Intensity) of PD-L1 were measured on the different monocyte subsets by flow cytometry. Results are presented as median with interquartile range. Statistical differences were determined by Kruskall-Wallis test (*) and Dunn's multiple comparisons test (#). ToD=Time of Diagnosis; EoT=End of Treatment; HNEC=healthy non-endemic controls. C=classical monocytes. I=intermediate monocytes. NC=non-classical monocytes.
