## Supplementary Figure for "Altered co-stimulatory and inhibitory receptors on monocyte subsets in patients with visceral leishmaniasis"

### Slide 1
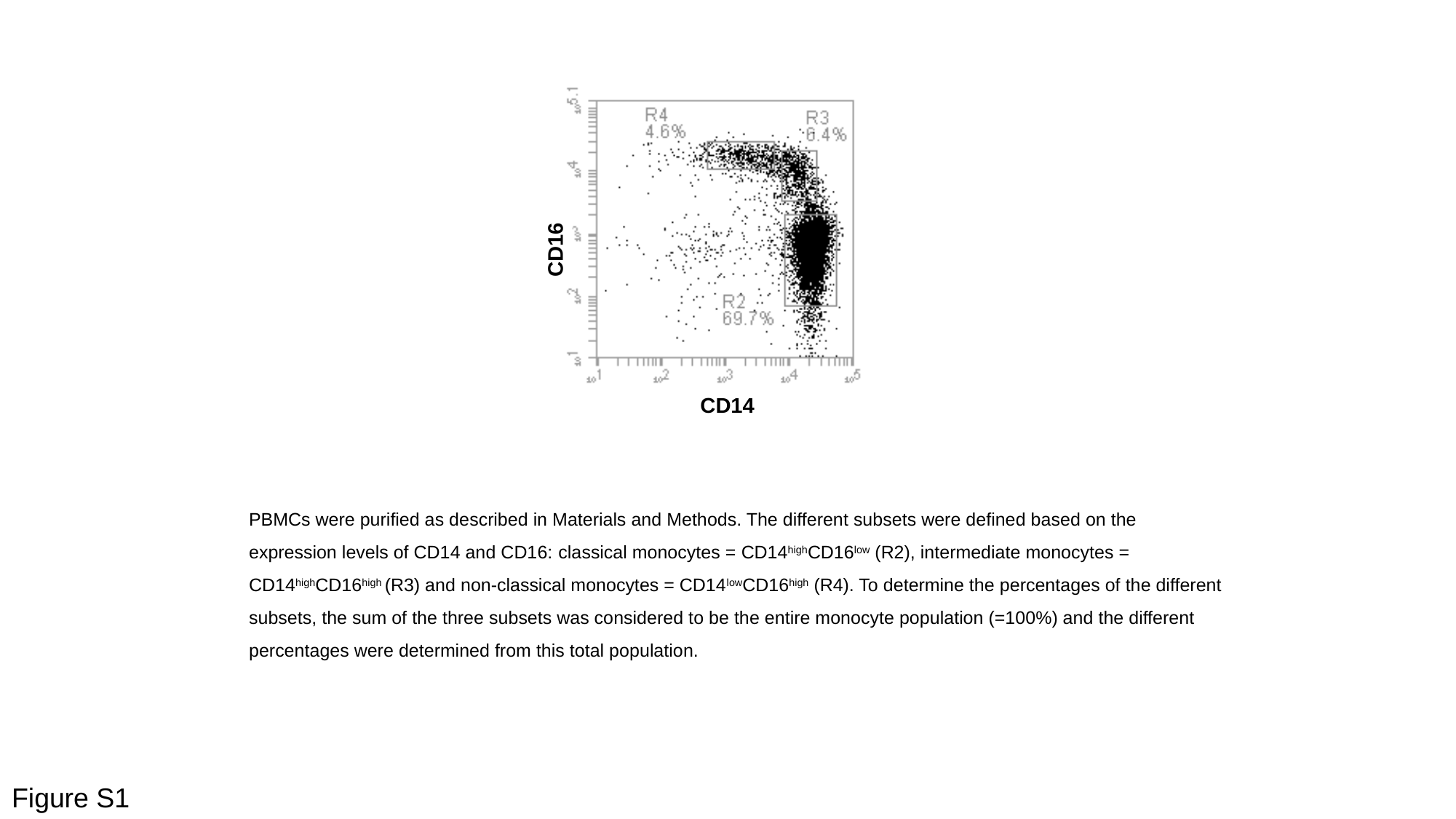

CD16
CD14
PBMCs were purified as described in Materials and Methods. The different subsets were defined based on the expression levels of CD14 and CD16: classical monocytes = CD14^high^CD16^low^ (R2), intermediate monocytes = CD14^high^CD16^high^ (R3) and non-classical monocytes = CD14^low^CD16^high^ (R4). To determine the percentages of the different subsets, the sum of the three subsets was considered to be the entire monocyte population (=100%) and the different percentages were determined from this total population.
Figure S1
